# Antagonism of kappa opioid receptors accelerates the development of L-DOPA-induced dyskinesia in a preclinical model of moderate dopamine depletion

**DOI:** 10.1101/2023.07.31.551112

**Authors:** Andrew J. Flores, Mitchell J. Bartlett, Grace Samtani, Blake T. Seaton, Morgan R. Sexauer, Nathan C. Weintraub, James R. Siegenthaler, Dong Lu, Michael L. Heien, Frank Porreca, Scott J. Sherman, Torsten Falk

## Abstract

Levels of the opioid peptide dynorphin, an endogenous ligand selective for kappa-opioid receptors (KORs), its mRNA and pro-peptide precursors are differentially dysregulated in Parkinson’s disease (PD) and following the development of L-DOPA-induced dyskinesia (LID). It remains unclear whether these alterations contribute to the pathophysiological mechanisms underlying PD motor impairment and the subsequent development of LID, or whether they are part of compensatory mechanisms. We sought to investigate nor-BNI, a KOR antagonist, 1) in the dopamine (DA)-depleted PD state, 2) during the development phase of LID, and 3) via measuring of tonic levels of striatal DA. While nor-BNI (3 mg/kg; *s.c.*) did not lead to functional restoration in the DA-depleted state, it affected the dose-dependent development of abnormal voluntary movements (AIMs) in response to escalating doses of L-DOPA in a rat PD model with a moderate striatal 6-hydroxdopamine (6-OHDA) lesion. We tested five escalating doses of L-DOPA (6, 12, 24, 48, 72 mg/kg; *i.p.*), and nor-BNI significantly increased the development of AIMs at the 12 and 24 mg/kg L-DOPA doses. However, after reaching the 72 mg/kg L-DOPA, AIMs were not significantly different between control and nor-BNI groups. In summary, while blocking KORs significantly increased the rate of development of LID induced by chronic, escalating doses of L-DOPA in a moderate-lesioned rat PD model, it did not contribute further once the overall severity of LID was established. While we observed an increase of tonic DA levels in the moderately lesioned dorsolateral striatum, there was no tonic DA change following administration of nor-BNI.

**Highlights:**

- Mild L-DOPA-induced dyskinesia develops in moderately lesioned parkinsonian rats
- In the moderately-lesioned dorsolateral striatum tonic dopamine is increased
- Antagonizing dynorphin does not affect parkinsonian motor symptoms in rodents
- Antagonizing dynorphin increases rate of development of L-DOPA-induced dyskinesia
- Tonic dopamine in dorsolateral striatum is unchanged after antagonizing dynorphin

## 1. Introduction

Alterations in the expression of endogenous opioid peptides and their precursors, and in the expression of opioid receptors, have previously been observed in Parkinson’s disease (PD) and levodopa (L-DOPA)-induced dyskinesia (LID). Gerfen and coauthors (Gerfen, 1992; Gerfen et al., 1990; Gerfen & Scott Young, 1988) found that striatonigral projecting medium spiny neurons (MSNs) predominantly express D1 dopamine receptors and substance P, while striatopallidal projection MSNs express predominantly D2 dopamine receptors and the opioid peptide enkephalin. MSNs also co-release opioid peptides; specifically, the D2 receptor-expressing indirect striatopallidal projection neurons co-express enkephalins (Leu- and Met-) derived from the precursor pre-proenkephalin A (PPE-A, Penk), which preferentially bind and activate δ-opioid receptors, whereas the D1 receptor-expressing direct striatonigral projection neurons, in addition to Substance P, co-express dynorphin. Dynorphin, in particular, is a ligand selective for kappa (κ)-opioid receptors (KORs) (Breslin et al., 1993; Cuello & Paxinos, 1978; Gerfen & Scott Young, 1988; Seizinger et al., 1984). Brotchie’s group further expanded the knowledge of contributions of striatal opioid signaling, including the role of KORs (Brotchie et al., 1993; Fox et al., 2008; Hill & Brotchie, 1995; Hughes et al., 1998; Maneuf et al., 1995) and also investigated non-subtype-specific opioid antagonism in a clinical trial that showed this may prove useful in the treatment of L-DOPA-related wearing-off in PD but not in dyskinesia (Fox et al., 2004). Cenci’s group has revealed that LID in the rat is associated with striatal overexpression of prodynorphin mRNA (Cenci et al., 1998) and that bilateral alterations in cortical and basal ganglia levels of opioid receptor transmission were also closely associated with LID in rats (Johansson et al., 2001; Westin et al., 2001). Subsequent studies showed that (Henry et al., 2001, 2003) striatal levels of dynorphin are positively correlated with LID (Sgroi et al., 2016). Following dopamine (DA) depletion, striatal levels of PPE-A mRNA and enkephalin are upregulated, whereas levels of prodynorphin (PPE-B, Pdyn) mRNA and dynorphin in the striatum are downregulated (Calon et al., 2002; Henry et al., 2003; Nisbet et al., 1995). Therefore, while there is abundant evidence that opioid peptide expression is altered in PD and LID, the nature of the role played by these alterations in the pathophysiology of these disorders is unknown.

Activation of KORs and G_i_-signaling inhibits the release of DA in various midbrain efferents, including the nucleus accumbens (NAc) and the ventral striatum (Di Chiara & Imperato, 1988; Gehrke et al., 2008). Consistent with this, it was shown that administration of a KOR antagonist increased DA release in the nucleus accumbens (Spanagel et al., 1992), yet the question of whether this same phenomenon holds true in the dorsolateral striatum has not been answered. If true, then blockade of KORs in the motor part of the striatum could have motor-restorative effects in PD or affect the development of LID by enhancing DA release. Indeed, effects of KOR modulation in preclinical models of LID and PD have been demonstrated, for example, one study showed that opioid signaling is overactive in LID (Sgroi et al., 2016). Therefore, blocking KORs may also affect the development of LID. With regard to expression of KORs, Aubert et al. (2007) found that they were unaltered in MPTP-lesioned primates, but that their expression was reduced in the globus pallidus exterior (GPe) and globus pallidus interior (GPi) following chronic L-DOPA administration. In contrast, KOR expression was decreased in the striatum and SN, but unaltered in the GP, of 6-OHDA-lesioned rats that expressed AIMs in response to L-DOPA compared with non-dyskinetic animals (Johansson et al., 2001). Furthermore, activation of KORs with the selective κ-receptor agonist U50,488 reduced AIMs in the 6-OHDA rat model of LID and dyskinesia in MPTP-lesioned squirrel monkeys but also reversed the anti-parkinsonian effect of L-DOPA worsening motor behavior (Cox et al., 2007).

Based on this evidence, we hypothesized that blocking KORs will increase tonic DA in the DLS in moderate PD. We evaluated the potential modulatory effects of endogenous KOR signaling using the KOR-selective antagonist nor-binaltorphimine (nor-BNI) in three experimental paradigms: 1) a functional motor restoration PD paradigm utilizing a moderate unilateral striatal 6-hydroxydopamine (6-OHDA) rat model to recapitulate early PD and to test for restoration of motor functionality, 2) a prolonged LID development paradigm relying on chronic, escalating doses of L-DOPA following a moderate striatal 6-OHDA lesion to test if nor-BNI either increases or attenuates the development of LID, and 3) we then directly evaluated if nor-BNI modulates tonic DA levels in the dorsolateral striatum (DLS) using *in vivo* fast-scan controlled adsorption voltammetry (FSCAV).

## 2. Results

### 2.1. Functional restoration study: Post hoc analyses to verify unilateral 6-OHDA lesion

In *post hoc* analyses for the functional restoration study, a significant reduction in nigral TH immunoreactivity (**Figure 2A**) was observed in both vehicle/control (*n* = 5) and nor-BNI (*n* = 4) groups relative to the intact hemisphere, verifying successful development of a unilateral lesion of the nigrostriatal tract (*F_[3, 14]_* = 25.92, two-tailed *p* < 0.0001, one-way ANOVA). Tukey’s multiple comparisons *post hoc* tests revealed significant reductions in TH-ir (vehicle intact vs. vehicle lesion: mean diff. = 380.5, q = 7.656, df = 14, two-tailed *p* < 0.0005; nor-BNI intact vs. nor-BNI lesion: mean diff. = 533.7, q = 9.605, df = 14, two-tailed *p* < 0.0001) as shown in **Figure 2B**. There was no significant difference in TH-ir between vehicle and nor-BNI intact hemispheres nor was there a significant difference between vehicle and nor-BNI lesioned hemispheres (vehicle intact vs. nor-BNI intact: mean diff. = - 157.1, q = 2.980, df = 14, two-tailed *p* = 0.1985; vehicle lesion vs. nor-BNI lesion: mean diff. = −3.870, q = 0.07341, df = 14, two-tailed *p* > 0.9999), indicating that the two groups were balanced in their levels of DA depletion prior to testing. Normalized TH-ir (% lesion/intact) was not significantly different between vehicle (mean = 28.086, SEM = 7.570) and nor-BNI (mean = 20.9075, SEM = 3.518) groups, indicating that animals were assigned to control and nor-BNI groups such that the two groups were balanced in their average level of DA denervation prior to testing (unpaired t-test, *t* = 0.7869, df = 7, two-tailed *p* = 0.4571). The same was true for the remainder of the animals in the vehicle/control (*n* = 4) and nor-BNI (*n* = 4) groups where analysis of DA, DOPAC, 5-HT, and 5-HIAA content in lysates sampled *post hoc* from striatal tissue revealed significant reduction in levels of DA (*F_[1,12]_* = 8.192, two-tailed *p* = 0.0146, two-way ANOVA) and DOPAC (*F_[1, 12]_* = 12.58, two-tailed *p* = 0.0040, two-way ANOVA) levels between the lesioned and intact hemispheres (**Figures 2C** and **D**), while 5-HT (*F_[1, 12]_* = 2.553, two-tailed *p* = 0.1361, two-way ANOVA) and 5-HIAA (*F_[1, 12]_* = 0.3154, two-tailed *p* = 0.5847, two-way ANOVA) levels remained unaltered by the 6-OHDA lesion as shown in **Figures 2E** and **F**.

**Figure 1.**
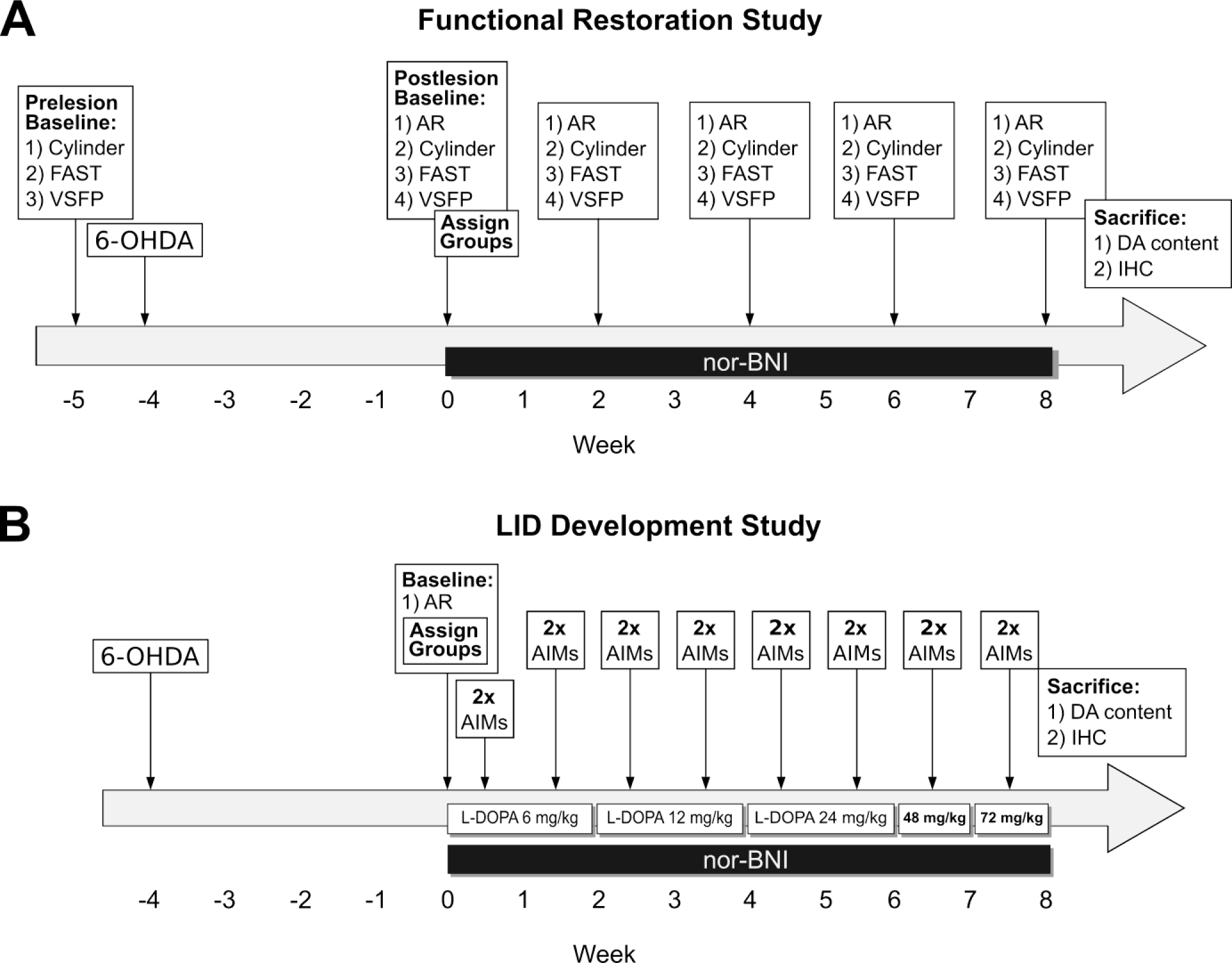
Experimental designs. **(A)** *Functional restoration paradigm:* We evaluated the potential of nor-BNI administration (3 mg/kg; s.c.) to restore motor functionality in a moderate unilateral striatal 6-OHDA lesion rat model (∼75% DA depletion). Within this paradigm, nor-BNI and vehicle-treated groups (*n* = 8-9) were tested multiple times using the amphetamine-induced rotations (AR) rotations test, the cylinder test, FAS test, and VSFP test to assess motor functionality. **(B)** *LID development paradigm:* We evaluated the effect of nor-BNI administration (3 mg/kg; s.c.) on the dose- and time-dependent development of abnormal involuntary movements (AIMs) in response to chronic escalating doses of L-DOPA treatment in a moderate unilateral striatal 6-OHDA lesion rat LID model (∼60% DA depletion). Within this mild-dyskinesia paradigm, independent nor-BNI and vehicle/control groups (*n* = 6-7) were tested multiple times (twice per week) across five escalating doses of L-DOPA (6, 12, 24, 48, and 72 mg/kg; *i.p.*) and evaluated for total limb axial and orolingual (LAO), and locomotor AIMs scores under blinded conditions.

**Figure 2.**
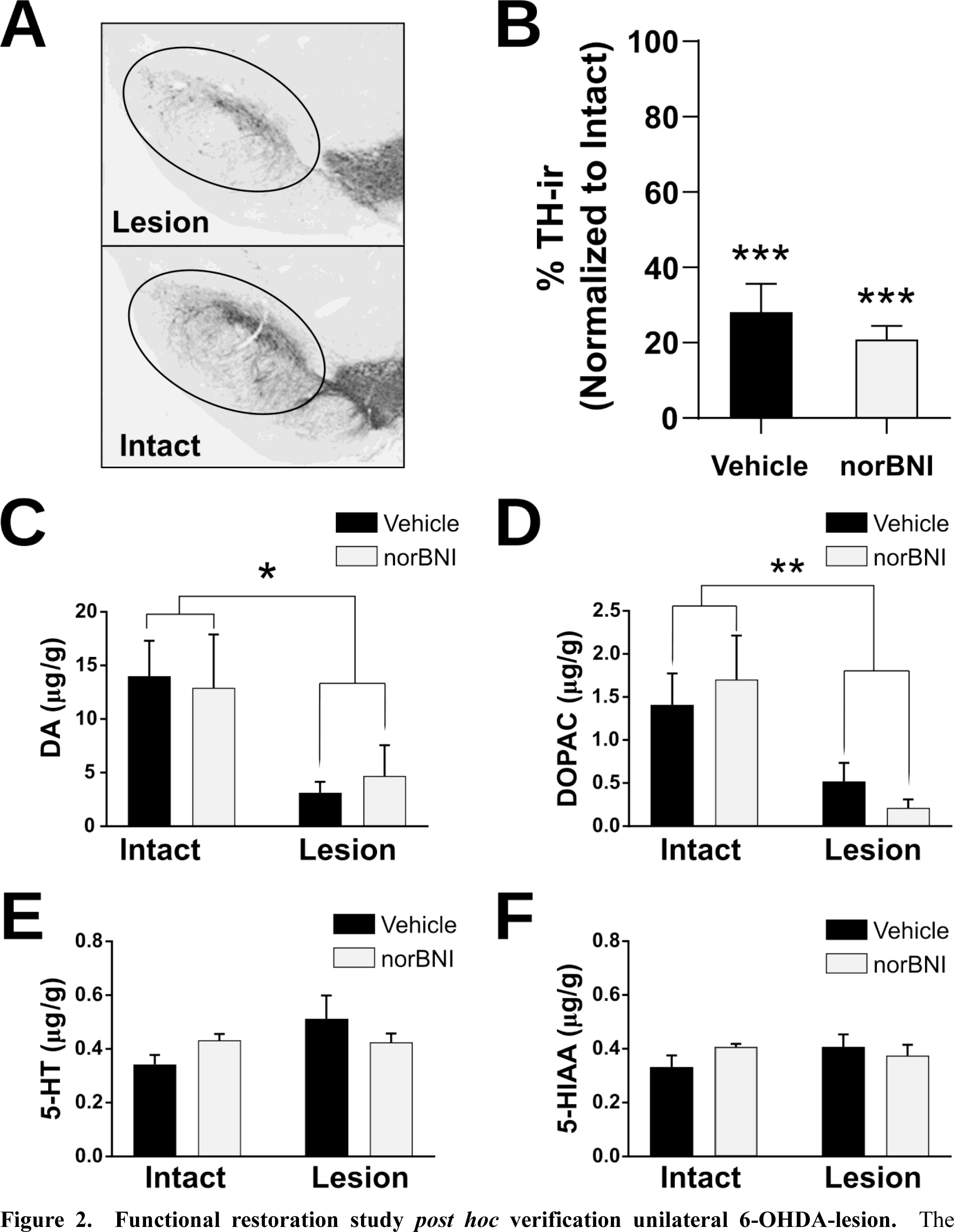
Functional restoration study *post hoc* verification unilateral 6-OHDA-lesion. The presence of unilateral lesions was verified using tyrosine hydroxylase (TH) immunoreactivity (TH-ir) as a marker of dopaminergic neurons in the substantia nigra for a subset of animals (**A** and **B**), and electrochemical measurement of striatal DA and DOPAC concentrations for the remaining animals **(C**-**F)**. The photomicrograph in **(A)** shows representative TH-ir in the substantia nigra (*upper panel:* 6-OHDA Lesion; *lower panel:* Intact) used for image analysis. The graph in **(B)** has plotted normalized TH-ir (% lesion/intact) and shows ∼75% reduction on the lesioned side (*n* = 4-5). In accordance with our mild unilateral 6-OHDA lesion paradigm, concentrations of striatal DA and its metabolite DOPAC measured with HPLC-EC (*n* = 4) were reduced by approximately 70 to 75% respectively in the lesioned striatum, while concentrations of serotonin (5-HT) and the serotonin metabolite 5-HIAA remained unaltered (**E** and **F**). Mean values ± SEM are plotted; * *p* < 0.05, ** *p* < 0.01, *** *p* < 0.001.

### 2.2. Treatment with nor-BNI did not improve motor deficits in the PD functional restoration paradigm

Analysis of the behavioral data from the AR, cylinder, FAS, and VSFP tests failed to detect any significant restorative effect of nor-BNI on parkinsonian motor deficits (**Figure 3**). For the AR, cylinder, FAS, and VSFP tests, two-way repeated measures ANOVA tests were used to compare the nor-BNI (*n* = 8-9) and vehicle/control (*n* = 9) groups across all post-lesion experimental timepoints. For the assays that had pre-lesion baseline scores, separate two-way ANOVA tests were performed to compare pre-lesion and post-lesion baseline scores, which, as expected, showed a marked reduction following the unilateral 6-OHDA lesion. Two-way repeated measures ANOVA was used to compare the average number of rotations per minute between control and nor-BNI groups in the AR test across all post-lesion timepoints (**Figure 3C**). This analysis revealed that there was no significant difference between control and nor-BNI groups in the FAS test (*F*_[4, 56]_ = 0.3432, two-tailed *p* = 0.8477). Two-way repeated measures ANOVA was used to compare forelimb touching bias scores from the cylinder test between vehicle/control (*n* = 8) and nor-BNI (*n* = 8) groups across all timepoints in the study. The analysis revealed that there was no significant difference in forelimb touching bias in the cylinder test between control and nor-BNI groups (*F*_[4, 56]_ = 0.8483, two-tailed *p* = 0.5007) as shown in **Figure 3B**. Two-way repeated measures ANOVA was used to compare the proportion of motor-defective forelimb steps to total steps taken in the FAS test across all post-lesion timepoints (**Figure 3C**). This analysis revealed that there was no significant difference between control and nor-BNI groups in the FAS test (*F*_[4, 56]_ = 0.8609, two-tailed *p* = 0.4931). Finally, a two-way repeated measures ANOVA was used to compare the proportion of reflexive touch responses elicited on the motor-defective left side to the intact right side across all post-lesion timepoints in the VSFP test (**Figure 3D**). This analysis revealed that there was no significant difference between control and nor-BNI groups in the VSFP test (*F*_[4, 56]_ = 1.456, two-tailed *p* = 0.2269).

**Figure 3.**
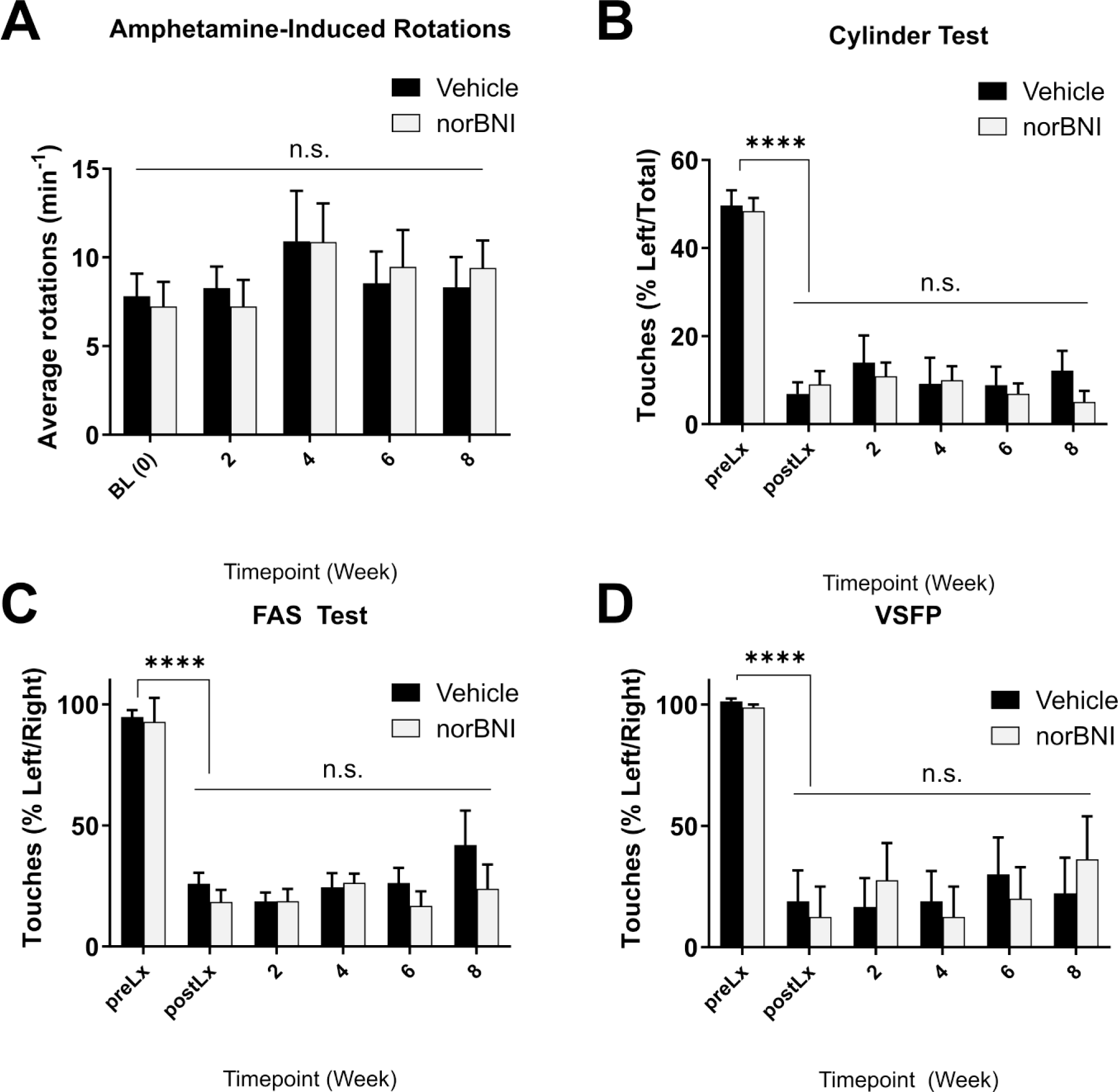
Functional restoration study: Behavioral analysis. **(A)** Amphetamine-induced rotations (AR) analysis failed to detect an effect of nor-BNI (3 mg/kg; *s.c.*) on the number of parkinsonian ipsilateral rotations per minute performed by the animals following amphetamine (5 mg/kg; *i.p.*) administration during timepoints following baseline (BL). Nor-BNI also did not show any motor-restorative effect in **(B)** FAS, **(C)** cylinder, nor **(D)** VSFP tests (preLx = prelesion; postLx = postlesion). Mean ± SEM, two-way repeated measures ANOVA, *n* = 8-9, **** *p* < 0.0005; n.s. = not significant.

### 2.3. LID development study: Post hoc analyses to verify unilateral 6-OHDA lesion

In the LID development study, *post hoc* HPLC-EC analysis was used to measure striatal concentrations of DA, DOPAC, 5-HT, and 5-HIAA in lysates derived from striatal tissue samples. Two-way ANOVA revealed that DA (**Figure 4A**) and DOPAC (**Figure 4B**) levels were significantly reduced in the lesioned versus the intact hemisphere (DA _lesion vs. intact_: *F*_[1,22]_ = 31.31, two-tailed *p* < 0.0001; DOPAC _lesion vs. intact_: *F*_[1,22]_ = 25.56, two-tailed *p* < 0.0001), thus verifying the successful development of the unilateral 6-OHDA lesion in animals, but that this reduction was not significantly different between nor-BNI and vehicle/control groups (DA _vehicle vs. nor-BNI_: *F*_[1,22]_ = 1.458, two-tailed *p* = 0.2400; DOPAC _vehicle vs. nor-BNI_: *F*_[1,22]_ = 9.876e-005, two-tailed *p* = 0.9922). This demonstrates that, as intended, animals were appropriately assigned to control and experimental groups, such that DA depletion was balanced between the two groups before commencing L-DOPA-therapy. By contrast, levels of 5-HT and 5-HIAA (**Figure 4C** and **4D**) between the lesioned and intact hemispheres were unaltered by the 6-OHDA lesion (5-HT: *F*_[1,22]_ = 0.001239, two-tailed *p* = 0.9722; 5-HIAA: *F*_[1,22]_ = 0.8918, two-tailed *p* = 0.3553) indicating that the lesion was specific to dopaminergic neurons.

**Figure 4.**
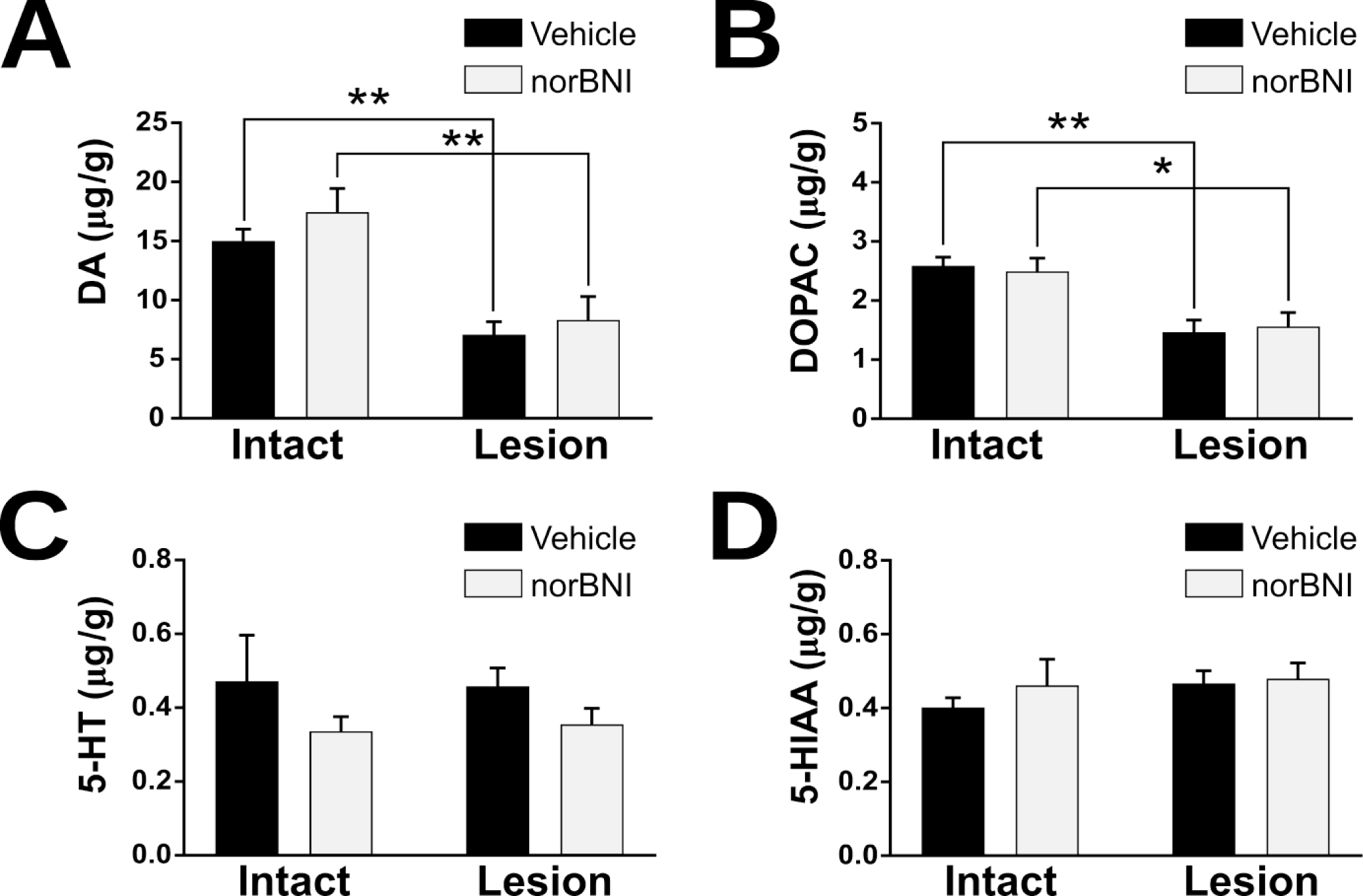
LID Development Study: *Post hoc* validation of extent of 6-OHDA-lesion. Electrochemical measurement of striatal DA concentrations was performed to verify levels of DA depletion associated with the unilateral 6-OHDA lesion. In accordance with our mild unilateral 6-OHDA lesion paradigm, concentrations of striatal DA (**A**) and its metabolite DOPAC (**B**) measured with HPLC-EC were reduced by approximately 50-60% in the lesioned striatum. Concentrations of serotonin (5-HT) shown in (**C**) and the serotonin metabolite 5-HIAA (**D**) remained unaltered. Mean values ± SEM are plotted, *n* = 6-7; * *p* < 0.05, ** *p* < 0.01.

### 2.4. Treatment with nor-BNI increased the rate of development of L-DOPA-induced AIMs in the model of prolonged LID development utilizing chronic escalating doses of L-DOPA

We evaluated the effect of *nor*-BNI administration (3 mg/kg; *s.c.*) on the dose- and time-dependent development of abnormal voluntary movements (AIMs) in response to chronic escalating doses of L-DOPA treatment in a moderate unilateral striatal 6-OHDA lesion rat LID model with ∼60% DA depletion (**Figure 1B**). The escalation paradigm was based on the standard dose escalation paradigm for development of dyskinesia in severely lesioned 6-OHDA rats with an escalation from 6 to 12 mg/kg after 2-weeks, established by Cenci’s group. We reasoned, given the moderate lesion we are using here (60-70% vs. >95% severe lesion), we would need higher doses of L-DOPA to induce AIMs. Therefore, within this mild-dyskinesia paradigm, independent nor*-*BNI and vehicle-treated groups (*n* = 6-7) were tested twice per week across 5 escalating dose phases of L-DOPA (6, 12, 24, 48, 72 mg/kg; *i.p*) and evaluated for total limb axial and orolingual (LAO; **Figure 5A**), and rotation/locomotor (**Figure 5B**) AIMs scores under blinded conditions.

**Figure 5.**
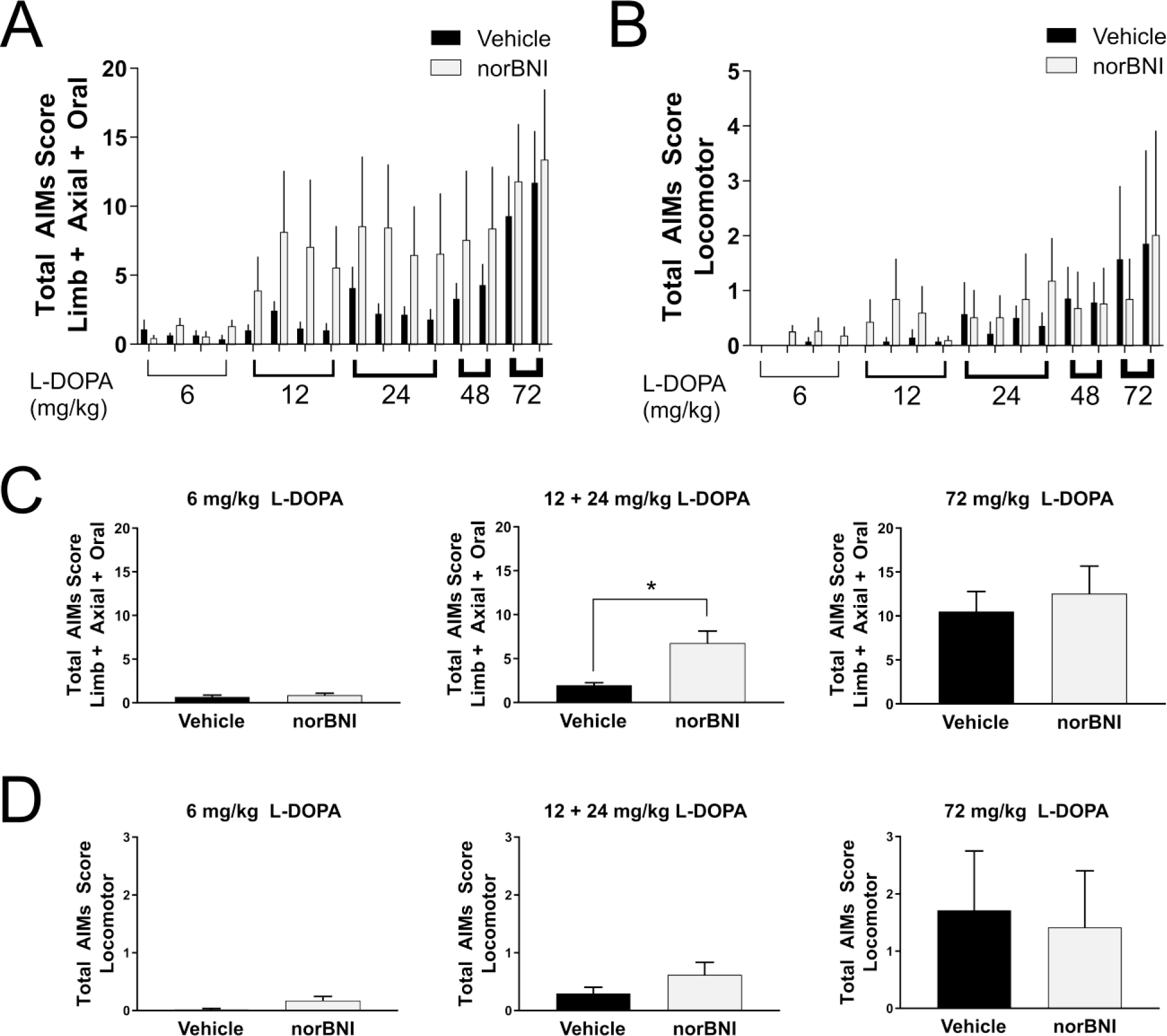
Treatment with nor-BNI increased the development of LAO and Locomotor AIMs. Animals were treated with gradually increasing doses of L-DOPA (6, 12, 24, 48, & 72 mg/kg). (**A**) Limb Axial Oral composite and (**B**) Locomotor AIMs scores obtained across the duration of the study, and a Kruskal-Wallis tests with Dunn’s *post hoc* tests revealed that there was no significant effect of nor-BNI on single treatment days. Bar plots indicate mean values ± SEM; *n* = 6-7. (**C**) Cumulative total LAO AIMs scores were analyzed separately with two-tailed Wilcoxon tests according to L-DOPA dose, and showed while there were no expression of LAO AIMs at 6 mg/kg L-DOPA, that nor-BNI increased the expression of LAO AIMs at the 12 mg/kg and 24 mg/kg L-DOPA doses during LID development by 240%. At the 72 mg/kg L-DOPA dose, LAO AIMs scores were not different between the two groups, suggesting that nor-BNI accelerates LID development, but does not change LAO AIMs magnitude once LID is established. (**D**) Cumulative total Locomotor AIMs scores were analyzed separately according to L-DOPA dose. Locomotor AIMs were not significantly different between groups at 6, 12 + 24, and 72 mg/kg L-DOPA doses. Bar plots indicate mean values ± SEM, * *p* < 0.05.

When cumulative AIMs scores were analyzed separately according to L-DOPA dose, we discovered that while no LAO AIMs were apparent at 6 mg/kg L-DOPA, nor-BNI significantly increased the expression of LAO AIMs (**Figure 5C**) by 240% at the 12 and 24 mg/kg L-DOPA doses (two-tailed Wilcoxon test, *p* = 0.0409). At the end of the escalation paradigm, after the 72 mg/kg L-DOPA dose, AIMs scores were the same in both groups (two-tailed Wilcoxon test, *p* = 0.9697), suggesting that nor-BNI accelerates the development of L-DOPA-induced AIMs, but does not increase the overall severity of AIMs once LID is established. When cumulative total rotation/locomotor AIMs scores were analyzed (**Figure 5D**) separately according to L-DOPA dose at the 6, 12 + 24 and 72 mg/kg doses locomotor AIMs were equal. In summary, nor-BNI significantly accelerated the development of LAO but not locomotor AIMs induced by chronic, escalating doses of L-DOPA in a moderate striatal lesion rat PD model, but did not increase either AIMs once LIDs were established.

### 2.5. Tonic DA is increased in the moderately lesioned DLS and treatment with nor-BNI did not alter tonic DA levels

Tonic levels of DA were measured in the DLS using FSCAV following administration of nor-BNI (3 mg/kg, *s.c.*) in anesthetized naïve animals and animals with a moderate striatal 6-OHDA lesion (**Figure 6A-D**). The lesioned animals selected for the study had an amphetamine-rotation count per minute of 3.8 ± 1.6 (mean ± SEM; *n* = 6), indicating a moderate lesion. A two-way repeated measures analysis of data recorded from animals in the vehicle/control (*n* = 5) and nor-BNI (*n* = 8) groups revealed that nor-BNI administration did not alter tonic DA levels (*F*_[1,11]_ = 0.06483, two-tailed *p* = 0.8037). Bonferroni *post hoc* tests yielded *p* > 0.05 for all timepoints following nor-BNI administration. In the moderate striatal 6-OHDA lesion group, tonic DA levels were binned every 10 minutes and a one-way repeated measures ANOVA was performed comparing the baseline, vehicle timepoints, and nor-BNI timepoints which showed nor-BNI had no effect on tonic DA in the striatum (*F*_[17,85]_ = 0.1740, two-tailed *p* > 0.9999). The implantation sites were verified *post hoc* (**Figure 6D**). Surprisingly, the tonic DA levels in the DLS were significantly higher in the 6-OHDA lesioned hemisphere when compared to the levels in naïve animals (**Figure 7A-C**). This ∼40% increased average tonic striatal DA in the lesioned DLS was significant for each of the three conditions (unpaired t-test, naive vs lesion baseline: t = 3.591, df = 17, two-tailed *p* = 0.0023; naive vs lesion vehicle: t = 2.947, df = 9, two-tailed *p* = 0.0163; naive vs lesion nor-BNI: t = 2.533, df = 12, two-tailed *p* = 0.0263). The moderate nature of the striatal lesion, was verified *post hoc* using western blot analysis of the SN (∼75% loss of DA cells) and near-infrared staining in the striatum (∼80% loss of DA terminals), after staining for tyrosine hydroxylase (TH), as shown in **Figure 8A-C**.

**Figure 6.**
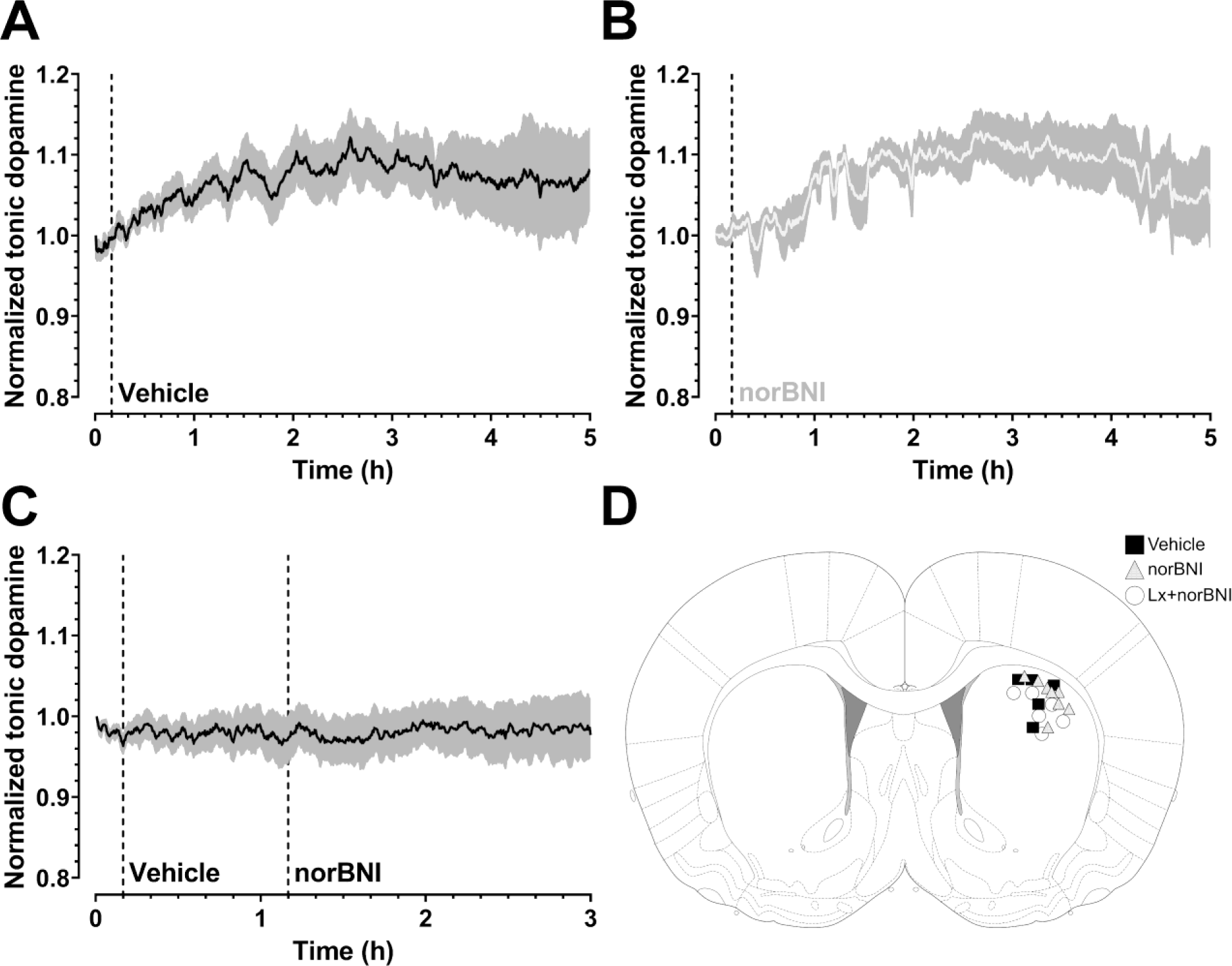
Subcutaneous injection of nor-BNI (3 mg/kg) does not alter tonic dopamine levels in the DLS. Normalized tonic DA levels, determined using FSCAV, after subcutaneous injection of (**A**) vehicle or (**B**) nor-BNI in naïve animals, and sequential injections of first vehicle and then nor-BNI in animals with a moderate striatal lesion (**C**). Mean ± SEM is plotted; *n* = 5 rats (vehicle/control), *n* = 8 rats (nor-BNI), *n* = 6 rats (lesioned); dotted lines represent injection time. A two-way repeated measures ANOVA showed no significant effect of subcutaneous nor-BNI on tonic DA levels in the DLS in the naïve animal group (*F*_1,11_ = 0.06483, two-tailed *p* = 0.8037). Bonferroni *post hoc* test: *p* > 0.05 for all timepoints. A one-way repeated measures ANOVA found that there was no significant effect of subcutaneous nor-BNI injection on tonic DA levels in the DLS for the lesioned animal group (*F*_[17,85]_ = 0.1740, two-tailed *p* > 0.9999). (**D**) The DLS recording locations included in dopamine analysis are shown in the scheme.

**Figure 7.**
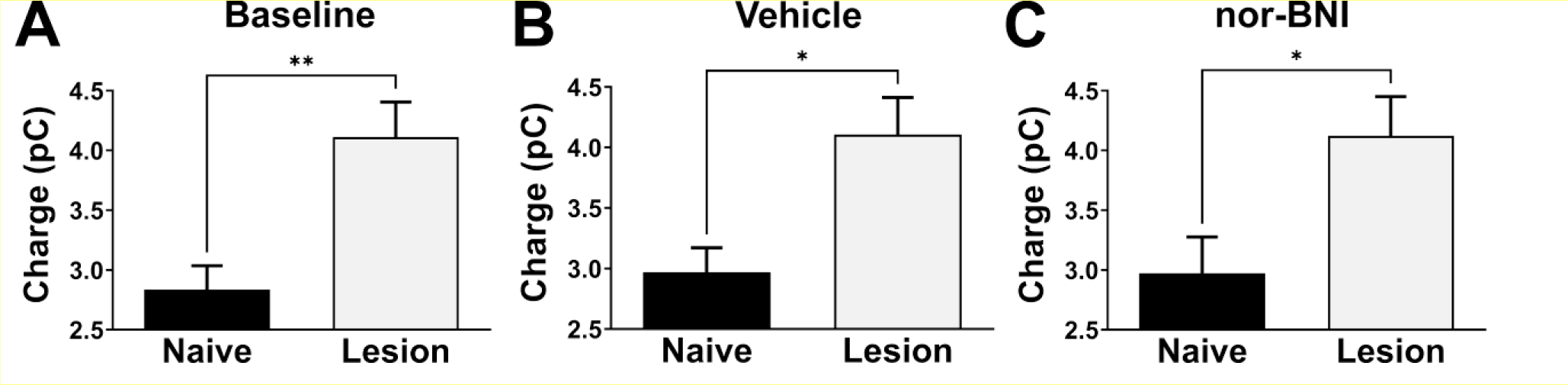
The tonic DA levels in the moderately lesion striatum are increased compared to the na. ï**ve animal levels.** Comparisons of average tonic DA levels between naïve animals with intact striatum (black bars) and animals with a moderate striatal lesion (grey bars) during (**A**) the baseline, (**B**) after vehicle administration, and (**C**) after nor-BNI administration. In each comparison, an unpaired *t*-test found that lesioned animals had significantly ∼40% higher tonic DA levels compared to the naive control animals (naive vs lesion baseline: *t* = 3.591, df = 17, two-tailed *p* = 0.0023; naive vs lesion vehicle: *t* = 2.947, df = 9, two-tailed *p* = 0.0163; naive vs lesion nor-BNI: *t* = 2.533, df = 12, two-tailed *p* = 0.0263). Mean ± SEM is plotted; *n* = 13 rats (naive baseline), *n* = 5 rats (naive vehicle), *n* = 8 rats (naive nor-BNI), *n* = 6 rats (lesioned); * *p* < 0.05, ** *p* < 0.01.

**Figure 8.**
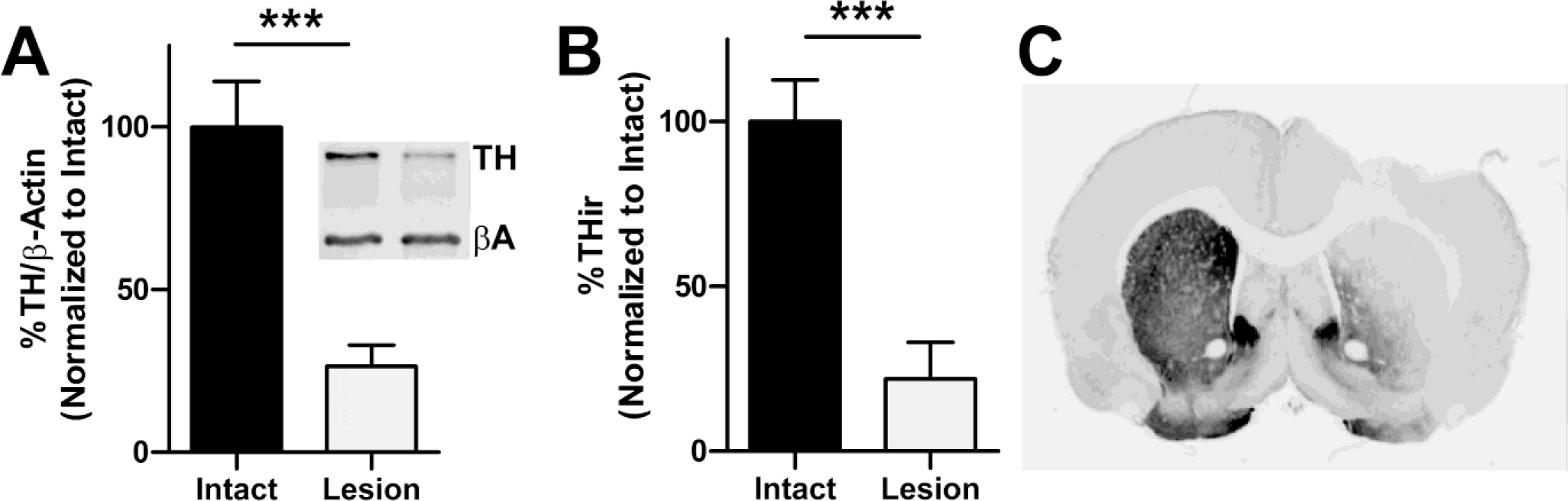
Verification of the moderate striatal lesion used in the voltammetry study. The unilateral 6-OHDA lesion used in Experiment 3, voltammetry measurement of tonic DA, was verified using tyrosine hydroxylase (TH) immunoreactivity (TH-ir) as a marker of dopaminergic cells and terminals. In **(A)** semi-quantitative western analysis of nigral tissue revealed ∼75% loss of DA cells. The inset shows an example blot (βA = beta actin). In **(B)** striatal TH-ir (normalized to % intact hemisphere) has been plotted and shows ∼80% reduction of terminals on the lesioned side. The photomicrograph in **(C)** shows TH-ir in a representative striatal section used for image analysis. Mean values ± SEM are plotted; two-tailed *t*-tests, *** *p* < 0.001; (*n* = 6).

## 3. Discussion

### 3.1. Blocking KOR signaling did not lead to motor restoration in moderate PD but accelerated development of LID

The opioid neurotransmission system is prominent in the basal ganglia. Endogenous opioid peptides and their precursors are highly expressed within the basal ganglia where they are modulators of GABA, DA, and glutamate neurotransmission (Fox et al., 2006; Fox & Brotchie, 2011). Several lines of evidence suggest that κ-mediated opioid signaling is overactive in LID. In the unilateral 6-OHDA lesioned rat model of LID, the severity of AIMs correlates strongly with levels of striatal PPE-B mRNA (i.e., prodynorphin), which encodes endogenous ligands for KORs (Cenci et al., 1998). Another study showed that striatal PPE-B mRNA levels were increased in both MPTP-lesioned macaques and PD patients with LID (Henry et al., 2003). Several other studies have shown that striatal levels of dynorphin are positively correlated with LID (Sgroi et al., 2016). This is in line with dynorphin levels being increased with many lesions in the brain and is not specific to this type of injury (Hauser et al., 2005), as well as a role for dynorphin/KOR signaling in brain circuits that are part of the stress response (Bruchas et al., 2010). Additionally, MALDI-TOF imaging mass spectrometry showed that levels of dynorphin B and alpha-neoendorphin in the SN are strongly correlated with the severity of AIMs in the 6-OHDA rat model of LID (Ljungdahl et al., 2011). Furthermore, [35S]GTPγS binding indicated hyperactive KOR signaling in the caudate nucleus and motor cortex in a nonhuman primate model of LID (L. Chen et al., 2005). Consistent with this concept, the relatively selective long-acting (Kishioka et al., 2013) KOR antagonist nor-BNI, has previously been shown to block L-DOPA-induced sensitization after nigral infusion in DA-depleted rats (Newman et al., 1997). By contrast, activation of KORs with the exogenous agonist TRK-820 has also been reported to reduce LID (Ikeda et al., 2009) and to increase locomotor activity in the monoamine-depleted rat model of parkinsonism (Hill & Brotchie, 1999; Hughes et al., 1998b). Despite this evidence, however, antagonizing KORs with nor-BNI failed to reduce levodopa-induced abnormal involuntary movements in primates with established LID (Henry et al., 2001), but this had not been investigated during the development of LID.

We found that nor-BNI increased the development of L-DOPA-induced AIMs at the 12 and 24 mg/kg doses by 240% using a paradigm of protracted gradual LID development in rats with a moderate 6-OHDA-lesion, modelling an earlier PD stage. Yet it did not worsen LID once it was fully established at 72 mg/kg, the latter replicating what has been shown in the primate model of established LID (Henry et al., 2001). In addition, nor-BNI also did not lead to any motor restoration in a model of moderate PD. While our initial hypothesis was that blocking KORs may increase tonic DA release in the DLS, as it does in the ventral striatum, one could also expect the opposite given the complex connectivity of basal ganglia circuitry. Direct striatonigral pathway dynorphin signaling also affects basal ganglia output neurons in the *substantia nigra pars reticulata* (SNr), downstream from the DA synapses in the MSNs. DA depletion produces a decrease in dynorphin expression in the direct pathway, which can be taken to reflect a decrease in activity in that pathway due to the loss of D1 receptor stimulation. According to classic rate models of basal ganglia function, loss of direct pathway activity that is inhibitory in nature contributes to overactivity of basal ganglia output and behavioral deficits. Hyperkinetic abnormal involuntary movements relate to DA receptor hyperstimulation (i.e., dyskinesia), in contrast to PD symptoms, that are thought to result from excessive indirect pathway activity driving thalamic inhibition, and concurrently, diminished direct pathway firing is thought to disinhibit output neurons further suppressing the thalamus (Graybiel et al., 2000). Blocking KORs could further decrease dynorphin/KOR signaling in the SNr, which might be expected to worsen lesion-induced deficits, not improve them. Given the lack of an effect in the motor restoration paradigm perhaps these two opposite mechanisms cancelled each other out.

The results from our studies are in contrast to the KOR effects in the NAc/ventral striatum, where it has been established that nor-BNI selectively blocks KOR signaling using multiple preclinical outcome measures including, for example, dysphoria (Land et al., 2008). Additionally, KOR agonists have been shown to induce dysphoric and aversive states (Xuei et al., 2006). Pain-induced negative affect is also mediated by activation of the kappa/dynorphin system in the NAc and in the central nucleus of the amygdala (Massaly et al., 2019; Navratilova et al., 2019). There has also been a great deal of work done testing the influence of KOR transmission associated with clinical depression, and within the context of withdrawal from opioid addiction and other substances of abuse (Chavkin & Koob, 2016; Knoll & Carlezon, 2010; Lalanne et al., 2014). One speculation on the differences in KOR-mediated modulation of tonic DA between DLS and ventral striatum could be the existence of different populations of KORs, perhaps differently saturated by endogenous dynorphin, affecting the nigral circuitry differently. Alternatively, differential neurotransmitter control of DA neural activity and corresponding DA release based on projection target has been shown, e.g. KOR agonists inhibit VTA DA neurons that project to the prefrontal cortex but not those that project to the NAc (Margolis et al., 2006). Nor-BNI could also act on the motor system outside the SN or even the basal ganglia, possibly via the habenula. KORs are expressed in the habenula and hyperexcitability of the lateral habenula is implicated in the development of major depressive disorder (Browne et al., 2018). Interestingly, while the lateral habenula is part of limbic systems involved in affective and emotional control, inhibiting the lateral habenula improves LID (Bastide et al., 2016; Cenci et al., 2018). The differential effects of KOR antagonism on tonic DA in motor and limbic systems might therefore depend on a complex combination of local but also system levels effects.

While we cannot rule out that dynorphin might be reflective of hyperactivity of the direct striatonigral pathway, our data further support compensatory involvement of KORs in LID, and indicate that KOR agonism could be an effective approach to therapy early on, during the beginning stages of the development of LID. On the other hand, the liability of KOR agonists is a concern, and likely will limit clinical use, specifically, in light of its role in driving aversion, negative affective states, stress-induced alterations in motivated behavior and cognition, as well as psychotomimetic effects (as reviewed in Tejeda & Bonci, 2019).

### 3.2. Considerations on the moderate lesion PD model used

Intrastriatal 6-OHDA administration induces damage of dopaminergic terminals, followed by a delayed, progressive loss of the dopaminergic neuron cell bodies in the SN, which are then secondarily affected through a dying-back phase. Considering this, we allowed 3-4 weeks for the striatal lesions to develop, to not interfere with our assessment of nor-BNI’s effects with this confounding variable.

Because terminals in the striatum are spatially diffuse relative to the densely bundled axons which travel between the SNc and striatum (included within the medial forebrain bundle tract), striatal administration of 6-OHDA causes a milder level of degeneration relative to more commonly-used lesions of the medial forebrain bundle, which cause severe (>95%) DA cell loss. For the functional Restoration study, we implemented a moderate lesion severity to realize ∼75% DA depletion, which is generally accepted as the level of cell loss required for motor deficits to appear in human PD patients. This leads to a sufficient level of motor dysfunction in order to measure any acute motor-restorative effects of nor-BNI, and the ∼25% remaining DA terminals would allow for any possible effects of nor-BNI on striatal DA release, as it has been shown for NAc and dorsomedial striatum (Di Chiara & Imperato, 1988; Gehrke et al., 2008). In the experimental paradigms to test LID development and the *in vivo* FSCAV, in addition to utilizing striatal lesions, we also used an even lower amount of 6-OHDA, to realize an even more moderate lesion of ∼60% DA depletion, to assure that a sufficient level of functional DA activity remained. This is different from the severe MFB lesions (>95% lesion), that model later PD stages and are the standard for the rat LID model, where development of LID can be induced by low L-DOPA doses, typically 6-7 mg/kg (Lundblad et al., 2002). Striatal lesions are typically not used for studies of LID because in this case very high doses of L-DOPA must be administered in order to induce AIMs. In paradigms using DA-depletion, L-DOPA doses as high as 50-100 mg/kg or more have been used to induce AIMs without toxicity (Engber et al., 1991; Ishida et al., 2000).

### 3.3. Tonic DA is increased in the DLS after a moderate 6-OHDA lesion and nor-BNI did not change tonic DA in the DLS of either naïve or hemi-lesioned PD animals

The increased rate of LID development in nor-BNI treated animals could provide evidence in support of the hypothesis that nor-BNI causes a functional increase in DA levels within the DLS, as a rise in tonic DA within the motor part of the striatum would act to correct the imbalance between the direct and indirect striatal output pathways, which fundamentally underlies PD pathology. Alternatively, opioid modulation of signaling downstream of DA could underlie the effects of nor-BNI to worsen the development of LID. Our FSCAV experiment in naïve and moderately 6-OHDA-lesioned rats showed that systemic nor-BNI did not change tonic DA levels in the DLS over the course of the 5-hour and 3-hour experiments, in contrast to the increased tonic DA levels shown by others in the ventral striatum. This argues against direct effects of nor-BNI on tonic DA levels in the DLS, but rather points to κ-opioid modulation of downstream intracellular signaling, due to the block of KORs, i.e. reduced activation of mitogen-activated protein kinases (MAPK), which should be explored in detail in future studies. In addition, our experiment did provide us with a surprising novel finding, that the tonic DA in the DLS after a moderate hemi-lesion (∼75%) did not decrease, but rather increases by 40% compared to naïve animals, pointing to the compensatory capacity of the DLS to make up for the loss of DA and possibly even overcompensate in a model of early stages of PD with only moderate dopaminergic cell loss. An increase in tonic DA after a moderate 6-OHDA lesion has not been investigated prior, though, Dave et al., 2014 found that Pink1 KO rats, a genetic PD model with a mild progressive loss of dopaminergic cells, had an increased DA content in the striatum compared to wild-type rats. Levels were elevated two- to threefold at eight months of age (Dave et al., 2014) before a later decline after 12 months. Young 3 month old G2019S mutant knock-in mice exhibit elevated DA transmission, before a decline at 12 months of age (Volta et al., 2017). Moreover, in mice overexpressing α-synuclein with a Thy-1 promoter elevated striatal tonic DA levels were seen at 6 months of age, prior to any reduction in total striatal tissue content (Lam et al., 2011). Knockdown of both α-synuclein and γ-synuclein was shown to lead to increased striatal DA release (Senior et al., 2008). Further, the pesticide dieldrin induced an increase in DA release in striatal slices in α-synuclein pre-formed fibril (PFF)-injected animals without changing vesicular monoamine transporter type 2 (VMAT2), suggesting that developmental dieldrin exposure increases a compensatory response to synucleinopathy-triggered striatal DA loss (Boyd et al., 2023). These data from different genetic PD models, where the disorder develops slowly over time, point to the possibility of overcompensation in tonic DA in the parkinsonian striatum in an earlier stage, a stage we are modeling with our moderate PD model using a rapid acting toxin. While we could not measure the striatal DA in tissue in Experiment 3, as the validation of the correct implantation site in the DLS was done with IHC, we do have the striatal DA tissue data from Experiment 1, where the same ∼75% nigral loss was achieved, leading to a ∼75% DA loss in the tissue. We can only speculate as to the mechanism involved in the ∼40% increase in tonic DA levels seen in tissue with ∼75% DA reduction, as the interplay among neuronal firing, DA release, and DA uptake is not only nonlinear and dynamic, but state- and region-dependent. Short repeated burst activity produces rapid presynaptic facilitation of DA release, while prolonged burst activity slowly depresses evoked DA release (Somayaji et al., 2020). The net tissue DA concentration at any point in time results from a dynamic equilibrium between release and uptake (Sulzer et al., 2016), and pharmacological dopamine transporter (DAT) uptake inhibition increases outside DA concentration by 300% (Carboni et al., 1989; Gonon & Buda, 1985). We would suggest that perhaps the DA release probability was increased, possibly due to increased short repeated nigral burst firing, while there was only a moderate downregulation of DAT in the moderate lesion, leading in sum to an increase in tonic DA release despite reduced DA levels. It will be important to evaluate the underlying mechanisms in future studies utilizing different lesion severities.

### 3.4. Considerations of recent literature on other opioid receptors in PD and LID

Other recent studies have explored modulation of opioid signaling, including κ-opioid signaling, rather with dually acting compounds or within a combination treatment, i.e. the mixed κ-opioid receptor agonist/μ-opioid receptor antagonist nalbuphine did reduce LID in dyskinetic non-human LID primates (Potts et al., 2015). Further, a recent study did show that the novel molecule DPI-289, a mixed δ-opioid agonist/μ-opioid receptor antagonist, was anti-dyskinetic in both rodent and non-human primate LID models (Johnston et al., 2018). In addition, the mixed δ/μ-opioid receptor agonist MMP-2200 blocked apomorphine-induced hyperkinesia in 6-OHDA-lesioned rodents (Yue et al., 2011), and mitigated the pro-parkinsonian activity of the *N*-methyl-D-aspartate (NMDA) receptor antagonist MK-801 without interfering with the anti-dyskinetic activity (Flores et al., 2018). While a MOR antagonist with low selectivity over DOR (15-fold) reduced LID (Koprich et al., 2011), two highly selective MOR antagonists did not (Bartlett et al., 2020b; Bezard et al., 2020), and rather a MOR agonist showed anti-dyskinetic activity (Bezard et al., 2020). In addition, agonists at the nociceptin opioid peptide (NOP) receptor, the fourth member of the opioid receptor family, attenuate dyskinesia as well (Marti et al., 2012; Mercatelli et al., 2020). Together this indicates that more than one opioid receptor modulation might be needed for a successful anti-LID strategy given the complexity of the basal ganglia circuitry and its multifaceted modulation by the opioid system.

### 3.5. Limitations and Conclusions

A limitation of this work is that the studies were conducted in male rats, future studies should evaluate female rats to test if there are any sex-specific effects. The FSCAV experiments were conducted in anesthetized animals, and future work could expand this to FSCAV recordings in behaving animals, especially given our surprising novel finding, that tonic DA in the DLS was increased despite a reduced tissue DA level. In conclusion, our data indicate that while blocking endogenous KOR signaling did neither improve PD motor dysfunction, nor had an effect on established LID in a preclinical rodent model, it accelerated the development of LID in a rodent model of moderate PD. This further indicates that dynorphin is not only protective against a variety of brain insults, as shown by others, but that KOR signaling might modulate dysregulated neurotransmission within the basal ganglia in the parkinsonian state treated with L-DOPA. While we cannot rule out that dynorphin might be just reflective of hyperactivity of the direct pathway neurons our findings argue for a possible compensatory activity of dynorphin.

## 4. Experimental procedure

### 4.1. Experimental Design

#### PD Restoration paradigm

The design of the PD restoration experimental paradigm is diagramed in **Figure 1A**. In this paradigm, we evaluated the effects of nor-BNI (3 mg/kg; *s.c.*) to restore motor functionality in a moderate unilateral striatal 6-OHDA lesion rat LID model (∼75% DA depletion). Within this mild DA depletion paradigm, motor functionality was assessed in independent nor-BNI and vehicle-treated groups (*n* = 9/group; one animal in the nor-BNI group died of unrelated causes in week 7 post-treatment during this long-term study and some behavioral analysis videos for one animal in the control group were lost) using 1) amphetamine-induced rotations (AR), 2) cylinder, 3) forelimb-adjusting steps (FAS), and 4) vibrissae-stimulated forelimb placement (VSFP) tests in order to assess motor functionality.

#### LID development paradigm

The design of the LID-development experimental paradigm is illustrated in the schematic diagram in **Figure 1B**. In this paradigm, we evaluated the effect of nor-BNI administration (3 mg/kg; *s.c.*) on the dose- and time-dependent development of abnormal voluntary movements (AIMs) in response to chronic escalating doses of L-DOPA treatment in a mild unilateral striatal 6-OHDA lesion rat LID model (∼60% DA depletion). Animals (*n* = 14) with ≥1.0 amphetamine-induced rotations were included in the study (mean = 5.25, SEM = 1.30). Within this paradigm of prolonged LID development, independent nor-BNI and vehicle-treated groups (*n* = 7/group; one animal in the control group died of unrelated causes during this long-term study) were tested multiple times (2 testing sessions per week) across 5 escalating doses of L-DOPA (6, 12, 24, 48, 72 mg/kg; *i.p.*), and evaluated for total limb, axial, and orolingual (LAO) as well as rotation/locomotor AIMs scores under blinded conditions.

#### Measuring tonic striatal DA levels

In a third experiment we evaluated tonic DA levels in the DLS in anesthetized naïve animals and animals with a moderate striatal lesion using *in vivo* FSCAV in response to systemic nor-BNI (3 mg/kg; *s*.*c*.) administration. For the naïve group (*n* = 13), tonic DA was recorded for 5 hours with systemic vehicle or nor-BNI (3 mg/kg; *s*.*c*.) being administered 10-minutes into the experiment. For the moderately-lesioned group (*n* = 6) tonic DA levels were recorded for 3 hours, during which, sequential systemic vehicle and nor-BNI (3 mg/kg; *s*.*c*.) injections were administered 10-minutes and 70-minutes into the recording, respectively.

### 4.2. Animals

Male Sprague-Dawley rats (∼225 g at arrival; Envigo RMC Inc., Indianapolis, IN), were used for Experiments 1 and 2. For the DA voltammetry measurement in Experiment 3 adult male Sprague-Dawley rats (258-310 g; Envigo) were housed two per cage. Animals from all studies were housed on a 12-hour light/dark cycle with food and water provided *ad libitum*. All procedures were approved by the Institutional Animal Care and Use Committee at the University of Arizona and conformed to the guidelines of the NIH Guidelines for the Care and Use of Laboratory Animals, and with ARRIVE guidelines. All attempts were made to minimize the total number of animals used in these studies and their suffering.

### 4.3. Stereotaxic surgery for 6-hydroxydopamine striatal lesions

Two separate doses of 6-OHDA were used to induce chronic mild striatal lesions in animals— one for the functional restoration study and another for the LID-development study. For the functional restoration paradigm, a total amount of 15.0 μg 6-OHDA per animal was delivered into the striatum. For the LID-development paradigm, a total amount of 12.0 μg 6-OHDA per animal was delivered into the striatum. Prior to surgery, animals (*n* = 30 for the restoration paradigm and *n* = 18 for the LID development paradigm and *n* = 6 for the FACAV experiment) were acclimated in the animal facility for approximately 2 weeks until they reached an average body mass of 300 g. In both experimental paradigms, a total volume of 8.0 μL of freshly prepared 6-hydroxydopamine hydrobromide solution was given (Sigma-Aldrich, St. Louis, MO); 1.875 and 1.5 μg per μL in 0.9 % sterile saline with 0.02% ascorbic acid to deliver in two injection sites in the DLS 7.5 μg/injection for the restoration and voltammetry studies, or 6.0 μg/injection for the LID-development and FSCAV study, respectively. 6-OHDA was injected into two locations in the striatum with the following coordinates (1) AP +1.6; ML +2.2; DV −5.0; and (2) AP −0.4; ML +4.0; DV −5.0, according to the atlas of Paxinos & Watson (2007) as in (Yue et al., 2014). Injections were carried out at a rate of 1.0 μL per min using a 10.0 μL syringe and needle (26-gauge, 22 mm, 45°tip; Hamilton Company, Reno, NV) with a microinjector (Stoelting Co., Wood Dale, IL) attached to a stereotaxic micromanipulator (Narishige, Tokyo, Japan); the needle was left in place post-injection for a further 5 min to prevent any backflow of 6-OHDA solution upon withdrawal of needle. Given the established, standardized nature of the unilateral 6-OHDA model, no sham surgery groups were included.

### 4.4. Amphetamine-Induced Rotations (AR) test

In the functional restoration study, amphetamine-induced rotations were used to assess the severity/extent of dopaminergic denervation and asymmetric DA release occurring between the lesioned and intact hemispheres of the brain following development of unilateral 6-OHDA striatal lesions. Using AR is the most sensitive behavioral measure to assign groups in 6-OHDA-lesioned animals based on severity of lesion, and can distinguish a moderate lesion reliably, which other tests such as cylinder or FAS tests cannot do reliably (Björklund & Dunnett, 2019). The low criterion of 1 was used to identify animals with a moderate lesion, as compared to the standard criterion of 4-5 that indicates a severe lesion. Unilaterally-lesioned rats were injected with D-amphetamine (5.0 mg/kg, *i.p.*; Sigma-Aldrich) to induce dopamine release from dopaminergic neurons projecting to the striatum and were then placed in a plexiglass cylinder (38 cm diameter × 38 cm height). Rotational behavior was recorded by a camera placed above the cylinder for a total of 100 minutes following administration of amphetamine. The average number of rotations per minute were counted and calculated by an individual blinded to the experimental condition, and trained to distinguish a true stereotypic rotation from exploratory behavior. Amphetamine washes out within two days, and the repeated tests in Experiment 1 were spaced by two weeks.

### 4.5. Cylinder test

Cylinder behavioral tests were conducted in plexiglass cylinders (14.5 cm diameter; 30.0 cm height). Forepaw touches on cylinder walls were counted by a blinded observer on a video for a minimum of 20 touches during a 5–10 minute recording period and calculated by a formula given in (Schallert, 2006): [(ipsi + 1/2 both)/(ipsi + contra + both)] × 100, which sets non-bias at 50%, and plotted as % contralateral forepaw contacts.

### 4.6. Forelimb-adjusting steps (FAS) test

The potential for restoration of motor functionality was assessed using a technique previously described (Chang et al., 1999; Eskow et al., 2007). Briefly, a 0.9-meter distance is marked off on a smooth tabletop surface. The animal is held by the experimenter with one hand supporting, but obstructing movement of the hindlimbs. The hind limbs are then slightly raised off the surface of the table. The experimenter’s second hand is used to support and obstruct the movement of the forelimb not being assessed. The forelimb being assessed is allowed to touch the table and then moved sideways over the set distance at a rate of 0.1 m/s. The experimenter then moves the forelimb first in the backhand direction, followed by the same motion in the forehand direction. Each forelimb was assessed three times per testing session. The total number of adjusting steps made in each direction during the three trials were recorded and later counted and used to calculate the average number of steps according to limb and direction (forehand vs. backhand). The proportion of the average number of steps taken by the motor defective left limb to the intact right limb for a given directionality (i.e., forehand vs. backhand) is then calculated. All tests were performed by a blinded observer. This test was utilized in both the LID-development and functional restoration paradigms.

### 4.7. Vibrissae-stimulated forepaw (VSFP) test

The VSFP test was used to assess sensorimotor deficits in parkinsonian animals of the functional restoration study (Schallert, 2006). Animals were held parallel to the floor, with their hindlimbs supported and one forelimb restrained. The vibrissae on the side of the unrestrained limb were stimulated against the corner of a table in a single, constant motion. This normally causes the unrestrained forelimb to respond by reflexively touching the tabletop, unless a motor deficit is present in that limb due to the unilateral 6-OHDA lesion. Ten trials were performed for each forelimb and the proportion of touches on the motor defective left side to touches on the non-affected right side were calculated.

### 4.8. L-DOPA-Induced AIMs

For the LID-development study, following development of a moderate striatal lesion, animals were treated with chronic escalating doses L-DOPA. Within this mild-dyskinesia paradigm, independent nor-BNI and vehicle-treated groups were tested multiple times (twice per week; **Figure 1B**) across five escalating doses of L-DOPA (6, 12, 24, 48, 72 mg/kg; *i.p.*) and evaluated for total limb axial and orolingual (LAO), and locomotor AIMs scores. All injections were administered with the dopa decarboxylase inhibitor benserazide (14 mg/kg; *i.p.*) to prevent peripheral conversion of L-DOPA to DA and to mimic clinical therapy of human PD patients; injections were given once daily throughout the duration of the study. AIMs were tested twice per week (3 days between each testing session).

AIMs were assessed by an observer blind to treatment condition, using an adapted version of the Abnormal Involuntary Movement Scale (Cenci et al., 1998), previously described by Paquette et al. (2010) and in our previous publications (Bartlett et al., 2016; Bartlett, Flores, et al., 2020; Flores et al., 2014, 2018). Briefly, each animal was rated on a scale of 0 (absent) to 4 (most severe) on each of four subscales (limb dyskinesia, axial dystonia, oral dyskinesia, and contraversive rotation). Limb, axial, oral, and locomotor type AIMs were based on 1-minute observation intervals conducted every 20 minutes for 180 minutes after administration of drugs. Specifically, for limb dyskinesia, a rating of 1 indicated 1 discrete period of abnormal movement, 2 indicated 3 or more discrete periods of abnormal movement, and 3-4 indicated continuous abnormal movement, with 4 indicating that AIMs could not be interrupted by a loud tap on the test cage. For axial dystonia, a rating of 1 indicated a contralateral bias in head orientation, 2 indicated a contralateral bias in head and upper body orientation, and 3-4 indicated a severe contralateral bias in head and upper body orientation (i.e., head above the tail in a four-paw stance), with 4 indicating loss of balance (i.e., falling). For oral dyskinesia, a rating of 1 indicated 2-3 bouts of mastication, 2 indicated more than 3 bouts of mastication, and 3-4 indicated continuous mastication, with 4 indicating the presence tongue protrusions. For L-DOPA-induced contraversive rotation, a rating of 1 indicated 2-3 contraversive turns, 2 indicated more than 3 contraversive turns, and 3-4 indicated continuous contraversive rotation, with a 4 indicating that these could not be interrupted by a loud tap on the cage. When ipsiversive rotation was observed, this was rated identically to contraversive rotation, except a negative score was provided to indicate contraversive directionality. Limb, axial, and oral subscale scores were summed to create a composite LAO AIMs score, while the rotation score, often also referred to as locomotor AIMs scores were considered separately from LAO scores (Lundblad et al., 2002). Total scores for AIMs, as well as for each subscale, were summed over the entire duration of the experiment (180 minutes).

### 4.9. Drugs and Administration Protocols

In each study (functional restoration and LID-development), there were two independent groups (vehicle/control and nor-BNI). Assignment of animals to vehicle vs. experimental groups was based on post-lesion baseline performance in the AR test and also the cylinder and FAS test with the intention to balance the baseline motor functionality as much as possible between the groups prior to the commencement of testing (see **Figure 1**) with the intention of balancing the groups equally according to their level of motor deficit, which is in turn correlated to the severity of nigrostriatal degeneration and DA depletion. L-DOPA (levodopa; L-3,4-dihydroxyphenylalanine methyl ester hydrochloride), benserazide hydrochloride (D,L-serine 2-(2,3,4-trihydroxybenzyl) hydrazide hydrochloride) were purchased from Sigma-Aldrich, St. Louis, MO. Nor-binaltorphimine (nor-BNI) was purchased from Tocris Bioscience (Minneapolis, MN). In both the LID development and functional restoration paradigms, nor-BNI was administered subcutaneously (*s.c.*) at a concentration of 3 mg/kg three times per week, a dose that leads to long-term KOR antagonism in rodents (Navratilova et al., 2019). In the LID-development paradigm, nor*-*BNI was initially administered three days in advance of the commencement of L-DOPA administration, and three times per week to ensure complete ongoing blockade of KORs prior to AIMs testing. Nor-BNI was administered simultaneously with L-DOPA on testing days, L-DOPA and *nor*-BNI were administered at the beginning (*t* = 0 min) of testing. For the FSCAV study, naïve animals were split into a nor-BNI and vehicle/control group while 6-OHDA lesioned animals were administered both vehicle and nor-BNI sequentially with all injections being administered subcutaneously.

### 4.10. Electrochemical Measurement of DA and 5-HT and Metabolites

Rat brains were placed in chilled Tris buffer (pH = 7.4, 15 mM Tris, 125 mM NaCl, 2.5 mM KCl, 2 mM CaCl_2_) and gently agitated for approximately 30 s to remove any contaminants and then placed in a chilled brain block. Coronal brain slices were then collected and a 2 mm steel biopsy punch was taken to sample tissue from the striatum. Samples from left and right hemispheres were collected and immediately flash frozen on an aluminum pan placed in liquid nitrogen −70 °C. Samples massed at 2.5 ± 0.5 mg, were then placed in 1.5 mL homogenization vials with 100 μL of 0.1 N HClO_4_ (*aq*), manually homogenized (15 strokes) using a disposable pestle and stored at −80 °C for up to 2 weeks prior to analysis. High performance liquid chromatography with electrochemical detection (HPLC-EC) was subsequently used to separate and quantify DA, 3,4-dihydroxyphenylacetic acid (DOPAC), serotonin, and 5-hydroxyindoleacetic acid (5-HIAA).

### 4.11. Sectioning and immunohistochemical staining (Functional restoration study and voltammetry)

Following behavioral analysis, half the animals underwent immunohistochemical analysis for dopaminergic loss, as described in detail in prior work (Flores et al., 2018; Yue et al., 2011, 2014). Animals underwent perfusion-fixation using cold 4% paraformaldehyde, brains were dissected, and further fixed overnight in 4% paraformaldehyde. 40-micron sections through the substantia nigra and the striatum were obtained using a microtome (Leica Biosystems Nussloch GmbH) and mounted on standard glass slides (Fisher Scientific). Next, the sections were stained with a rabbit anti-TH primary antibody (1:10,000 for 24 hours at 4°C; Millipore) and a biotinylated donkey anti-rabbit immunoglobulin G secondary antibody (1:1,000 for 2 hours at RT; Millipore). To eliminate nonspecific binding prior to immunostaining endogenous peroxidase activity was blocked using 0.3% H_2_O_2_ for 30 min at room temperature (RT), the slides were then washed 3 times for 5 min with Tris buffer (pH = 7.6) and submerged in blocking solution (1% normal donkey serum, 0.1% Triton-X-100, Tris-HCl buffer, pH 7.6) for 2 hours at RT. The signal from the secondary antibody was amplified using the ABC reagent (Vectastain Elite ABC Kit, PK-6100) according to the manufacturer’s protocol. We used DAB substrate (Vector Laboratories, Burlingame, CA) as the chromogen for final visualization of TH immunoreactivity (TH-ir). Densitometry analysis of the total area of detected objects (total pixels^2^) in a 480 × 480 pixel region of the mediolateral SN (3 sections/rat) was conducted by a blinded investigator with ImageJ software (vs.2.0.0-rc-69/1.52p, http://rsbweb.nih.gov/ij/) according to an established protocol (Bartlett et al., 2020a; Kneynsberg et al., 2016).

Confirmation of the 6-OHDA lesion in the rats analyzed in Experiment 3 (DA voltammetry measurement) were confirmed using near-infrared imaging. One coronal section, containing the striatum, was selected at each of the following coordinates: AP +1.6, +1.0, −0.4 mm, relative to bregma (Paxinos & Watson, 2007). Sections were incubated in a blocking buffer containing Odyssey Intercept® (TBS) Blocking buffer (LI-COR Biosciences, Lincoln, NE) plus 10% normal donkey serum (Jackson ImmunoResearch Laboratories Inc., West Grove, PA) at room temperature for one hour. Sections were then incubated in blocking buffer, 0.1% Tween-20, and rabbit anti-TH hydroxylase primary antibody (1:2000; AB152, Millipore) overnight at 4 °C. Tissue was then washed 3 x 5 min in TBS + 0.1% Tween-20 (TBST) and incubated in blocking buffer and 0.1% Tween-20 including the IRDye® 800 CW Donkey anti-Rabbit (1:500; LI-COR Biosciences, Billerica, MA) secondary antibody for one hour at room temperature. Sections were washed again 3 x 5 minutes in TBST, mounted on subbed slides and imaged on an Odyssey CLx imager (800 nm channel, 42 μm resolution, 1.0 mm depth; LI-COR Biosciences). Quantification of TH-ir in the striatum was analyzed using Image Studio Lite version 5.2.5 based upon methods established in the literature with the following changes (Benskey et al., 2018; Gombash et al., 2013). TH-ir densitometry was quantified using Image Studio Lite software 5.2.5 (LI-COR Biosciences). Signal intensity was calculated in both the intact and lesioned hemispheres subtracting the background (cortex) intensity from the total signal of the hand traced striatum, which was then quantified as signal intensity per unit area (Signal intensity = (Total Signal – Background Signal)/Area). Signal intensity was then normalized to the intact striatum.

### 4.12. Semi-quantitative western analysis (voltammetry)

Tissue preparation and western blot analysis was performed as previously published in Bartlett et al., 2020, with the following changes. The tissue was dissected from both the intact and lesioned SN. A linear range for TH in the SN was determined and 4 μg of homogenate was loaded on to each gel, transferred, and stained with rabbit anti-TH (1:2,000, Millipore) and mouse anti-Beta-Actin (1:10,000, Sigma-Aldrich). Imaging and analysis were performed using an Odyssey CLx (LI-COR Biosciences) imaging system and analyzed with Empiria Studio Software (LI-COR Biosciences).

### 4.13. Fast-Scan Controlled Adsorption Voltammetry (FSCAV)

Electrodes: Carbon-fiber microelectrodes (CFMEs) were fabricated as previously described, with minor modifications (Gee et al., 2020). A single carbon fiber (7.1 μm diameter, AS4 12K, Hexcel, Stamford, CT) was aspirated into a glass capillary and pulled with a vertical micropipette puller (PE-2 Narishige, Tokyo, Japan). The exposed carbon fiber was cut to a length of 70 μm with a scalpel under a Micromaster brightfield microscope (12−561B, Fisher Scientific, Hampton, NH). A silver wire (Kauffman Engineering, Cornelius, OR), coated with alcohol-based graphite conductive adhesive (42465, Alfa Aesar, Ward Hill, MA), was inserted into the back end of the glass capillary. Epoxy adhesive (1373425, Henkel Corp., Rocky Hill, CT) was applied to the wire-capillary interface and allowed to dry overnight. Each CFME was then coated with 3,4-ethylenedioxythiophene:Nafion (PEDOT:Nafion) to increase DA sensitivity and selectivity (Vreeland et al., 2015). Ag/AgCl-wire reference electrodes were prepared by soaking 0.25 mm-diameter silver wire (265578, Sigma-Aldrich, St. Louis, MO) in 8.25% sodium hypochlorite (The Clorox Company, Oakland, CA) for 24 hours.

Surgery: Rats were induced with 4% isoflurane gas and an air flow rate of 1.5 L/min. Rat body temperature was maintained with a water-circulating heating pad (T/Pump, Stryker, Portage, MI). A bipolar stimulating electrode (Plastics One, Roanoke, VA) was implanted in the medial forebrain bundle (AP: −2.5 mm, ML: 1.7 mm, DV: −7.0 to −7.9 mm from bregma), a Ag/AgCl-wire reference electrode was implanted in the contralateral hemisphere, and a CFME working electrode was implanted in the dorsolateral striatum (DLS) (AP: 1.0 mm from bregma, ML: 3.2 mm from bregma, DV: −3.8 to −4.0 mm from brain surface). The final dorsoventral depth of the CFME was determined by maximizing stimulated DA release in the DLS. Following final electrode placement, tonic DA levels in the DLS were measured using fast-scan controlled adsorption voltammetry (FSCAV) (Atcherley et al., 2013). FSCAV was performed until a stable baseline was established (typically 30-60 minutes). For the naïve group, following baseline stability, FSCAV was performed for 10 minutes prior to injection of vehicle or nor-BNI (3 mg/kg) and continued for the remainder of the 5-hour experiment. For the lesioned group, following baseline stability, FSCAV was performed for 10 minutes prior to injection of vehicle and continued for 1 hour, followed by an injection of nor-BNI (3 mg/kg) and continued for the remainder of the 3-hour experiment.

Histology: Placement of the CFME working electrode was confirmed according to Gee et al., 2020 with the addition of cresyl violet staining. Following experimentation, electrolytic lesions were made with the working electrode and the animals were euthanized via isoflurane overdose and cervical dislocation. Brains were immediately post-fixed in 10% formalin and sectioned at 40 microns through the DLS on a cryostat (CM1850, Leica Biosystems, Buffalo Grove, IL). Sections were then stained with cresyl violet to improve visualization. The CFME was located within the DLS in each experiment, therefore all animals were included in the analysis. Representative histological verification is shown in **Figure 6D**.

Data processing: After collection, outliers were removed from the FSCAV data using the generalized extreme Studentized deviate test. For the naïve animal group, data were then normalized to the starting value for each animal and smoothed using a moving average with an 80-second centered window (the first two and last two data points had smaller windows of 20-second and 40-seconds respectively). For the lesioned group, outliers were removed, and the data were smoothed as above, however, instead of the data being normalized to the first data point, individual animal data was binned every 10 minutes for a total of 18 bins. These groups were then normalized to the 10-minute baseline bin for each animal. For the comparison between lesioned and intact DA levels, outliers were removed, and data were smoothed as previously mentioned, however, the data were not normalized.

### 4.14. Statistical Analysis

All statistical analyses were performed with GraphPad Prism 8 (GraphPad Software, La Jolla, CA). The TH-ir analysis was done using a one-way ANOVA with Tukey’s multiple comparisons *post hoc* tests. Two-way ANOVA was used to compare striatal DA, DOPA, 5-HT, and 5-HIAA content in lesioned and intact hemispheres of nor-BNI and vehicle/control animals. For the AR, cylinder, FAS, and VSFP tests, two-way repeated measures ANOVA tests were used to compare the nor-BNI and vehicle/control groups across all postlesion experimental timepoints; separate two-way ANOVA tests were used to compare prelesion and postlesion baseline scores. Within the LID development paradigm, non-parametric Kruskal-Wallis tests with Dunn’s multiple comparison *post hoc* tests were used to compare AIMs scores between nor-BNI and vehicle control groups across all experimental timepoints. Cumulative LAO and locomotor AIMs scores for the L-DOPA concentrations were compared between control and nor-BNI groups using non-parametric two-tailed Wilcoxon tests. For the FSCAV analysis, a repeated measures two-way ANOVA with Bonferroni’s multiple comparisons *post hoc* tests was used to compare the naïve nor-BNI and vehicle/control groups. For the lesioned animal group, a repeated measures one-way ANOVA with Tukey’s multiple comparisons *post hoc* tests was used. Two-tailed unpaired t-tests were used to compare the non-normalized average DA levels between the naïve and lesioned groups during baseline, after vehicle administration, and after nor-BNI administration. For all statistical analyses, two-tailed tests were performed to compare null and alternative hypotheses and the alternative hypothesis was rejected when *p* < 0.05. All graphs have plotted the sample mean ± SEM (std. error of mean).

## Acknowledgments

We would like to thank the ARCS Foundation (AJF), Training grant funding NIH T35 HL007479 (MJB) and NIH T32 GM008804 (NCW), and The Jerry T. and Glenda G. Jackson Fellowship in Parkinson’s Research to the University of Arizona (TF and SJS) for their support.

## CRediT author statement

**Andrew J. Flores:** Investigation, Visualization, Writing-Original draft preparation, Revision. **Mitchell J. Bartlett:** Investigation, Writing-Reviewing and Editing. **Blake T. Seaton:** Investigation, Writing-Reviewing and Editing. **Grace Samtani:** Investigation. **Morgan R. Sexauer:** Investigation. **Nathan C. Weintraub:** Investigation, Writing-Reviewing and Editing. **James R. Siegenthaler:** Investigation. **Dong Lu:** Investigation. **Michael L. Heien:** Supervision, Writing-Reviewing and Editing. **Frank Porreca:** Conceptualization, Resources, Writing-Reviewing and Editing. **Scott J. Sherman:** Writing-Reviewing and Editing, Funding acquisition. **Torsten Falk:** Conceptualization, Supervision, Writing-Reviewing and Editing, Revision, Resources, Funding acquisition.

